# Meta-analysis using new methods for three-stressor combinations reveal substantial higher-order interactions and emergent properties

**DOI:** 10.1101/2022.04.15.488520

**Authors:** Eleanor S. Diamant, Sada Boyd, Natalie Ann Lozano-Huntelman, Vivien Enriquez, Alexis R. Kim, Van M. Savage, Pamela J. Yeh

## Abstract

Although natural populations are typically subjected to multiple stressors, most past research has focused on single stressors and two-stressor interactions, with little attention paid to higher-order interactions among three or more stressors. However, higher-order interactions increasingly appear to be widespread. Consequently, we used a recently introduced and improved framework to re-analyze higher-order ecological interactions. We conducted a literature review of the last 100 years (1920-2020) and reanalyzed 151 ecological three-stressor interactions from 45 published papers. We found that 89% (*n=*134) of the three-stressor combinations resulted in new or different interactions than previously reported. We also found substantial levels of emergent properties— interactions that are only revealed when all three stressors are present. Antagonism was the most prevalent net interaction whereas synergy was the most prevalent emergent interaction. Understanding multiple stressor interactions is crucial for fundamental questions in ecology and also has implications for conservation biology and population management.

## Introduction

Individuals in natural populations almost always face multiple stressors that affect their ability to survive and to find food, shelter, mates, and safety (Blaustein & Kiesecker 2002; Côté *et al*. 2016). These stressors include changes to biological or environmental factors that can result in unfavorable responses within a population (Vinebrooke *et al*. 2004), leading to unfavorable responses across ecological systems (Jackson *et al*. 2016, 2021). Over the past century, ecological stressors such as climate change, pollution, and habitat destruction have adversely affected natural systems, contributing to biodiversity loss and a continuing threat to populations and ecosystems (Didham *et al*. 2007; Butchart *et al*. 2010; Halpern *et al*. 2015). These stressors rarely occur in isolation. Instead, they often interact—potentially changing the overall impact on populations (Crain *et al*. 2008; Côté *et al*. 2016). Therefore, there is a great need to properly assess and predict stressor interactions to mitigate their cumulative effects.

When the combined impact of two stressors is equal to the amount of the individual effects in isolation, the interaction type is defined as an additive interaction, or no interaction (Bliss 1939; Loewe 1953; Folt *et al*. 1999; Yeh *et al*. 2006; Piggott *et al*. 2015; Jackson *et al*. 2016). Alternatively, two stressors could interact synergistically—increasing the overall effects—or antagonistically—decreasing the overall effects (Bliss 1939; Loewe 1953; Folt *et al*. 1999; Yeh *et al*. 2006; Piggott *et al*. 2015). For example, synergistic interactions were observed when the combined effects of high temperatures and low pH decreased calcification (the production of shells and plates) in certain marine animals when compared to the individual effects of each stressor (Rodolfo-Metalpa *et al*. 2011). On the other hand, antagonistic interactions were found in coral (*Pocillopora meandrina*) microbiome response to multiple stressor interactions (Maher *et al*. 2019). Specifically, increased temperature and coral scarring both decreased the abundance of the dominant taxon (*Endozoicimonacae*) in coral microbiomes. However, the combined stressor effect led to a lower magnitude response than predicted if there were no interactions between the stressors (Maher *et al*. 2019). An extreme form of antagonism is termed suppression—one stressor reverses another stressor’s effects (Yeh *et al*. 2006; Chait *et al*. 2007; Singh & Yeh 2017). For example, the combined effect of carbaryl and nitrate decreased green frog (*Rana clamitans*) tadpole growth despite their individual positive effects on tadpole growth (Boone *et al*. 2005).

Pairwise interactions–the effects of two stressors in combination compared to individual effects– have been well studied in the ecological literature. Empirical work on pairwise stressor interactions (e.g., Hesse *et al*. 2012; Cramp *et al*. 2014; Van Praet *et al*. 2014; Sniegula *et al*. 2017; Delnat *et al*. 2019), literature reviews, and meta-analyses (Crain *et al*., 2008; Darling & Cote, 2008; Ban *et al*., 2014; Piggott *et al*., 2015; Jackson *et al*., 2016; Matthaei & Lange, 2016; Côté *et al*., 2016; Villar-Argaiz *et al*., 2018; Tekin *et al*., 2020) have revealed their substantial influence across biological systems and scales. Yet, there are likely more than two stressors acting on all, or almost all, wild populations. In fact, multiple stressor interactions are more frequent than previously thought, despite having received less attention (Beppler *et al*. 2016; Tekin *et al*. 2018a).

When studying interactions, higher-order combinations—defined here as a combination of three or more stressors—have long been ignored (Pomerantz 1981) despite their importance in ecological communities (Billick & Case 1994; Levine *et al*. 2017). The assumptions that have been used to justify this include (1) paired interactions or single-stressor effects provide the main effects, so one only needs to worry about paired interactions or single effects; higher-order interactions, therefore, provide negligible effects (Pomerantz 1981; Wootton 1994; Ban & Alder 2008; Wood *et al*. 2012; Wood 2016); (2) higher-order interactions are complex and depend on accurate, specific parameters and underlying null models that are often not available for effective and reliable testing (Billick & Case 1994; Thompson *et al*. 2018); and, (3) from an experimental standpoint, the collection of higher-order interactions, whether in the lab or the field, can be onerous, time-consuming, and logistically difficult as increasing stressor combinations could theoretically lead to an exponentially large number of experiments, which has given rise to research on approximating multi-stressor responses from single pairs (Billick & Case 1994; Côté *et al*. 2016; Wood 2016; Zimmer *et al*. 2016; though see: Levine *et al*. 2017).

A recently introduced framework to examine interactions was used specifically to analyze both pairwise and higher-order interactions (Tekin *et al*. 2018b, 2020). This framework, the Rescaled Bliss Independence (RBI), was originally drawn from the pharmacology and microbiology fields (Beppler *et al*. 2016; Tekin *et al*. 2016). The RBI has several key advantages compared to the most-used method to examine interactions in ecology, ANOVA. First, ANOVA incorporates several assumptions that are often violated or not tested (Text S1). Second, RBI enables direct comparisons of interaction effects from absolute to relative fitness. Third, the framework allows for straightforward generalization from pairwise to higher-order interactions while keeping the ability to rescale interaction terms—that is, to normalize interaction values relative to a natural baseline, much like the way we typically measure fitness as relative fitness rather than absolute fitness (see Text S2 for details). Finally, and crucially, the framework enables the identification of emergent properties—that is, what interactions arise that are the result of all three stressors together, rather than just being the dominant effect due to a pairwise interaction dominating the landscape of multiple stressors.

Emergent properties only arise from three or more stressors; there are no emergent properties in two-stressor interactions because the interaction between the two stressors is what emerges from the combination of the stressors. However, in three or more stressor combinations, the interaction from three stressors could be coming primarily from a strong two-stressor interaction—a non-emergent interaction—or the interaction could be the result of all three stressors together—an emergent interaction (Beppler *et al*. 2016). Together, pairwise and emergent interactions constitute the observed net three-way interaction (Figure 1A). For example, imagine ecological stressors A, B, and C all are impacting the growth of a population X. Let’s say that the interaction among all three stressors together is synergistic. But this three-stressor interaction could arise primarily from a strong synergistic pairwise interaction between two of the three stressors, say A and B, which would obscure all other interactions (described in Figure 1B). This would be a case of a non-emergent interaction. Or this three-stressor interaction could need all three stressors present to show a synergistic interaction, which would be an emergent interaction (Beppler *et al*. 2016; Tekin *et al*. 2016). Thus, it is imperative to quantify both pairwise and three-stressor combinations to determine the nature of population X’s response to A, B, and C.

**Figure 1.**
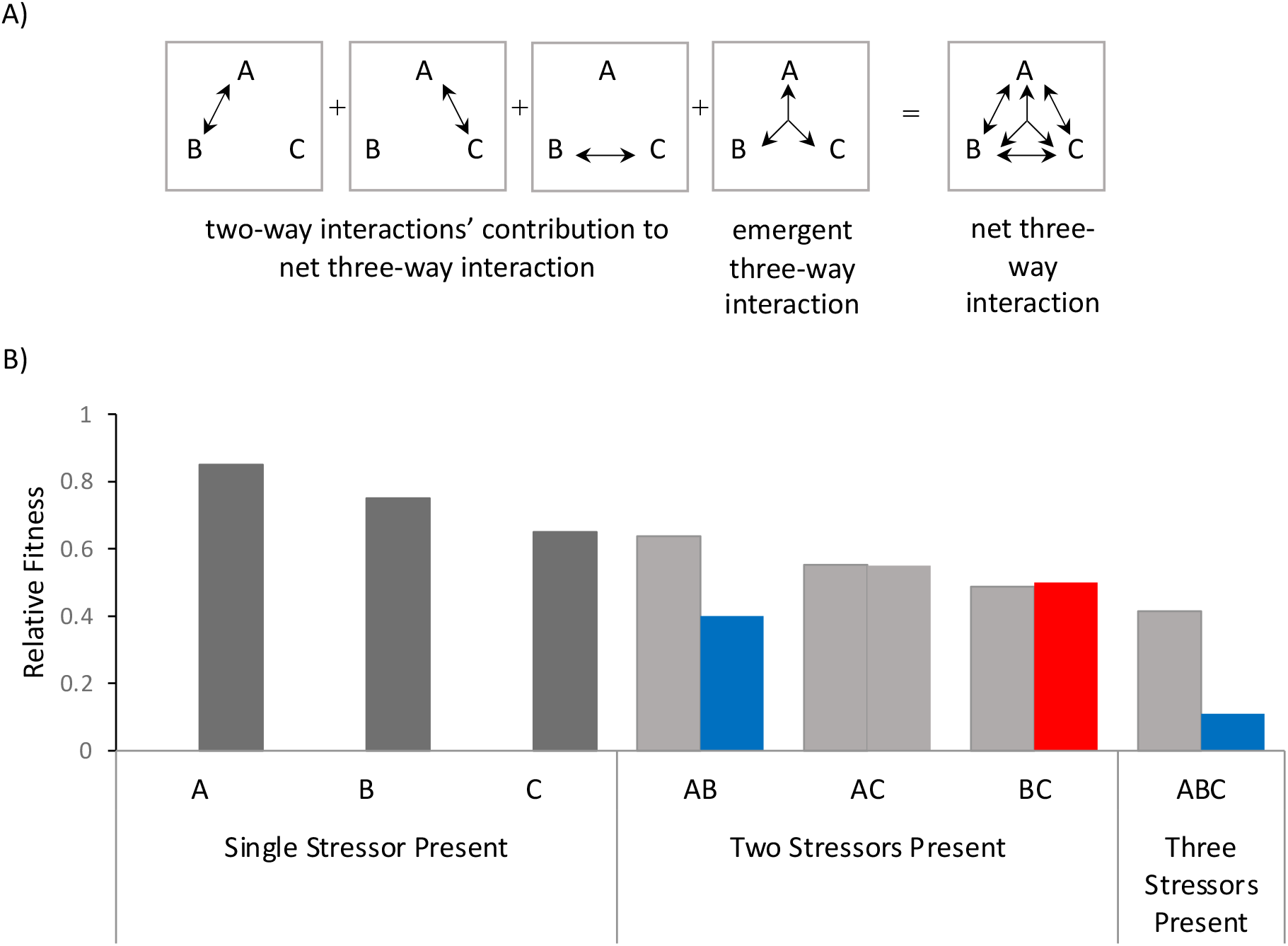
The components needed to assess three-stressor interactions. A) Net three-way interactions are composed of lower-order two-way interactions between given stressors (A, B, and C) and the higher-order emergent three-way interaction that is only quantifiable with all three stressors present and when all two-way interactions are known. Together, these compose the net three-way interaction. Figure 1 is partially adapted from (Tekin *et al*. 2018b). B) An example of the contribution of two-way interactions on net three-way interactions. Here, single stressors all decrease relative fitness. In two-stressor and three-stressor interactions, light gray represents the additive expectation based on single-stressor effects on fitness. Blue represents synergistic interactions and red represents antagonistic interactions. In this example, the strong synergistic two-stressor interaction between A and B, rather than an emergent interaction, overshadows the additive interaction between A and C and the slightly antagonistic interaction between B and C. This leads to a synergistic net three-way interaction between A, B, and C.

The importance of identifying emergent properties lies in our ability to understand the impacts of specific stressors, even when most populations experience multiple stressors in combination. In fields such as conservation ecology or climate change biology, there is often an emphasis on conserving and bolstering endangered and threatened populations by mitigating at least one of the stressors affecting population survival and growth (Brown *et al*. 2013). However, if we do not clearly understand how stressors interact, we could be mitigating the wrong stressors, or at least, not the optimal stressors. A striking example of the importance of emergent interactions can be seen in the field of pharmacology, where the combination of three antibiotic stressors trimethoprim, streptomycin, and erythromycin have a synergistic effect, efficiently reducing bacterial population size. However, if one of the drug stressors (for example, erythromycin) is removed from this combination the overall killing efficiency actually *increases* (Beppler *et al*. 2017). This results in the population of concern, the bacteria, decreasing more when there are only two stressors, rather than three (Beppler *et al*. 2017). This is an example of a critical emergent interaction. In much the same way, understanding multiple stressors and their emergent, higher-order, effects could be crucial for understanding how best to conserve species and populations.

Unlike ANOVA and similar approaches, the Rescaled Bliss Independence framework is conducive to rescaling the interaction measure (e.g., normalization), which allows for more easily measuring and identifying the strength of an interaction. Similar to absolute versus relative fitness, a rescaled interaction value is normalized by a natural baseline (e.g., lethality). Rescaling results in a multimodal distribution with clearer cut-offs around values. Therefore, interaction types are more easily distinguishable (Segrè *et al*. 2005; Tekin *et al*. 2016, 2020). Without the rescaling step, raw interaction values can be exactly the same even though they may represent different interaction types, leading to incorrect interpretations (Tekin *et al*. 2016). Therefore, rescaling is critical to comparing interaction measures and identifying interaction types.

Here we conduct a literature search of the last 100 years (January 1920-November 2020) and re-analyze stressor interactions using the new Rescaled Bliss Independence (RBI) framework (Tekin *et al*. 2016), recently applied to ecological studies to identify two-stressor interactions (Tekin *et al*. 2020). For simplicity, we define “stressors” as factors that affect population growth or fitness. While most of these “stressors” decrease population growth or fitness, a few stressors in the studies examined here actually increase population growth and/or fitness. We aim to obtain a more detailed, accurate, and complete understanding of higher-order ecological stressor interactions. Specifically, we use this framework to reanalyze the data from previously published papers (that used traditional methods i.e., ANOVA, General Linear Model, or log-logistic) that measure three-stressor interactions and all the lower-order interactions: all three pairwise combinations of stressors and all three single-stressors. We ask three questions: (1) How well does this new framework match previously published interaction results? (2) How often do emergent properties appear in higher-order ecological interactions? (3) Can we find patterns of emergent properties— that is, for example, do they primarily occur in synergistic interactions or antagonistic interactions?

## Materials and Methods

### Study Selection and Criteria

We conducted a literature search using the *Web of Science* database to select the studies included in our analysis. We searched one hundred years of published literature, from January 1920-November 2020, using the following key terms included in the papers’ keywords, title, and/or abstract: “multiple stressors,” “multiple antagonism,” “multiple synergy,” “multiple disturbance,” “multifactor,” “additions,” “indirect interactions,” and “stressors” (Supplemental Figure S1). Then, we further filtered the search results by selecting the following specific topic categories to reflect our interest in ecological studies: agriculture dairy, animal science, biodiversity conservation, biology, biotechnology applied microbiology, ecology, environmental sciences, evolutionary biology, genetics heredity, marine freshwater biology, microbiology, and zoology. We removed duplicate studies and only selected papers that measured growth, mortality, and/or survival at the population level for a specific species. Next, we examined the remaining papers to determine the presence of the following criteria: the study had (1) three individual stressors and a full multi-factorial design was implemented, (2) quantitative response variables, and (3) explicit control treatments. From the remaining papers, we extracted growth, mortality, and survival data from figures and tables from each of the qualifying studies. We included the following: stressor type, stressor units, responses for individual stressors, responses for combinations of stressors, responses for control variables, sample size, species of the organism tested, species natural habitat, and the interaction type between stressors determined by the original authors (i.e., additive, synergistic, or antagonistic). If a study did not specify if there was an interaction among the three stressors, we determined whether the authors specifically sought to investigate an interaction. Additionally, if a study reported that there was no interaction among the stressors—but the authors explicitly sought to investigate an interaction—we classified the interaction as additive because additivity is the null hypothesis when testing for interactions. Most of the quantitative responses from each study reflected mean values generated from raw values by the authors, often summarized from tables or figures provided in the studies. Other quantitative responses were directly obtained from raw data. We recorded the latest time point as the response value if mean or raw data were presented as a time series. Importantly, these factors could enhance or inhibit growth.

Finally, we filtered out combinations that would not work with the RBI framework. Specifically, RBI only works with uniform factors (e.g., all inhibitors or all enhancers of growth). Therefore, combinations that affected populations in opposite directions (e.g., a stressor decreasing population size and one increasing population size) were removed. Combinations that had a positive control value of zero could not be analyzed and were removed from the dataset. Combinations that had a lethal single stressor or a single stressor with no effect were assessed separately. In instances where one or more of the single stressors was lethal, we could only accurately identify the presence of a net suppressive interaction, and not an emergent effect. In this case, distinguishing additive, synergistic, or antagonistic interactions is not possible since a population cannot exhibit growth or survival less than zero. However, if the population demonstrates growth or survival in the presence of the three stressors combined, we could determine a suppressive interaction when at least one of the individual stressors is lethal. Additionally, there were cases where a single stressor had no effect. This can be problematic because it is unclear if the single stressor in combination with additional stressors has any effect or if we only see the effects of the additional stressors. Since we can only identify synergistic combinations under these circumstances, those cases were not included in our analysis. Many of the papers compared multiple combinations of three stressors. In total, 151 stressor combinations from 45 papers met the requirements needed for our study (Table S1).

### Data analysis

The RBI framework has previously been used to examine drug interactions and pairwise stressor interactions by relying on Bliss Independence as the additive model to determine if there is an interaction between stressors on a population (Beppler *et al*. 2016; Tekin *et al*. 2016, 2018a, 2020). We applied this framework to ecological studies exploring the impact of three stressors in each environment. Within this framework, there are a total of seven possible measurements one can take among the three stressors (stressor A, stressor B, and stressor C) acting simultaneously. These are: (1) the effects of A alone, (2) the effects of B alone, (3) the effects of C alone, (4) the pairwise effects of A and B by themselves, (5) the pairwise effects of A and C by themselves, (6) the pairwise effects of B and C by themselves, and (7) the effects of all three stressor A, B, and C together. The net interaction—termed deviation from additivity (*DA*) (Equation 1)—occurs when we remove the effects of the individual stressors from consideration. Removing the result of the pairwise interactions produces the emergent effect (*E3*) (Equation 2). Further, we can rewrite Equation 2 to only reflect relative fitness effects (Equation 3).

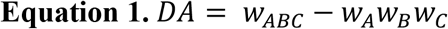

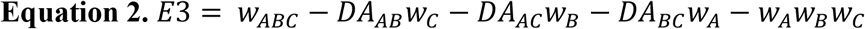

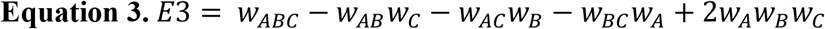

Upon calculating these initial interactions, rescaling methods and cutoff values used by Tekin *et al*. (2018b) were used to further investigate and identify interactions. After rescaling (see Text S2), both net and emergent interaction values below -0.5 were synergistic, values between -0.5 and 0.5 were additive, and values above 0.5 were considered antagonistic (where values above 1.3 were considered suppressive). For more information about rescaling and details on how to determine interactions for combinations where all stressors increase growth, see Text S2 and S3.

The framework described above requires that all single stressors have a non-lethal effect on relative fitness (0 < *w* ≠ 1). This is because if a single stressor is lethal (*w* = 0) or if a single stressor has no effect (*w* = 1) we would not be able to identify all interaction types. For example, if the use of one stressor results in complete lethality one cannot determine if a combination with that stressor interacts synergistically or additively if the combination also results in complete lethality. Similarly, if a single stressor appears to have no effect (*w* = 1) there is no way to distinguish if that stressor interacts at all with the system and is relevant or if the other stressor in the combination acts additively. This framework also requires relative fitness to be calculated with reference to a positive control (the growth of the population under no stressor present).

## Results

For our analysis, we collected data from multi-stressor ecological studies published over the last 100 years. Most of these studies were published in the past 10 years (Figure S2). We subsequently applied the RBI framework to reanalyze three-stressor interaction data derived from those studies. Our findings were then compared to those from the original studies. We found new net interactions that were previously unidentified by the original authors and net interactions that we reclassified based on our methods (Figure 2). Of the 151 interactions, nearly half, 42% (*n*=64), were interactions that were previously untested (e.g., experiments were conducted but no statistical analysis on the interactions themselves were reported) but classified as an interaction using RBI. We classified 17% (*n*=25) of previously tested but unspecified interactions (e.g., an interaction was found using statistical analyses by previous authors though the type of interaction was not explicitly stated). Only 11% (*n*=17) of the total interactions reanalyzed by RBI were re-classified with the same interaction type previously reported in the original studies while 30% (*n*=45) were different from what was previously reported. Collectively, 89% of interactions were newly classified or reclassified by RBI.

**Figure 2.**
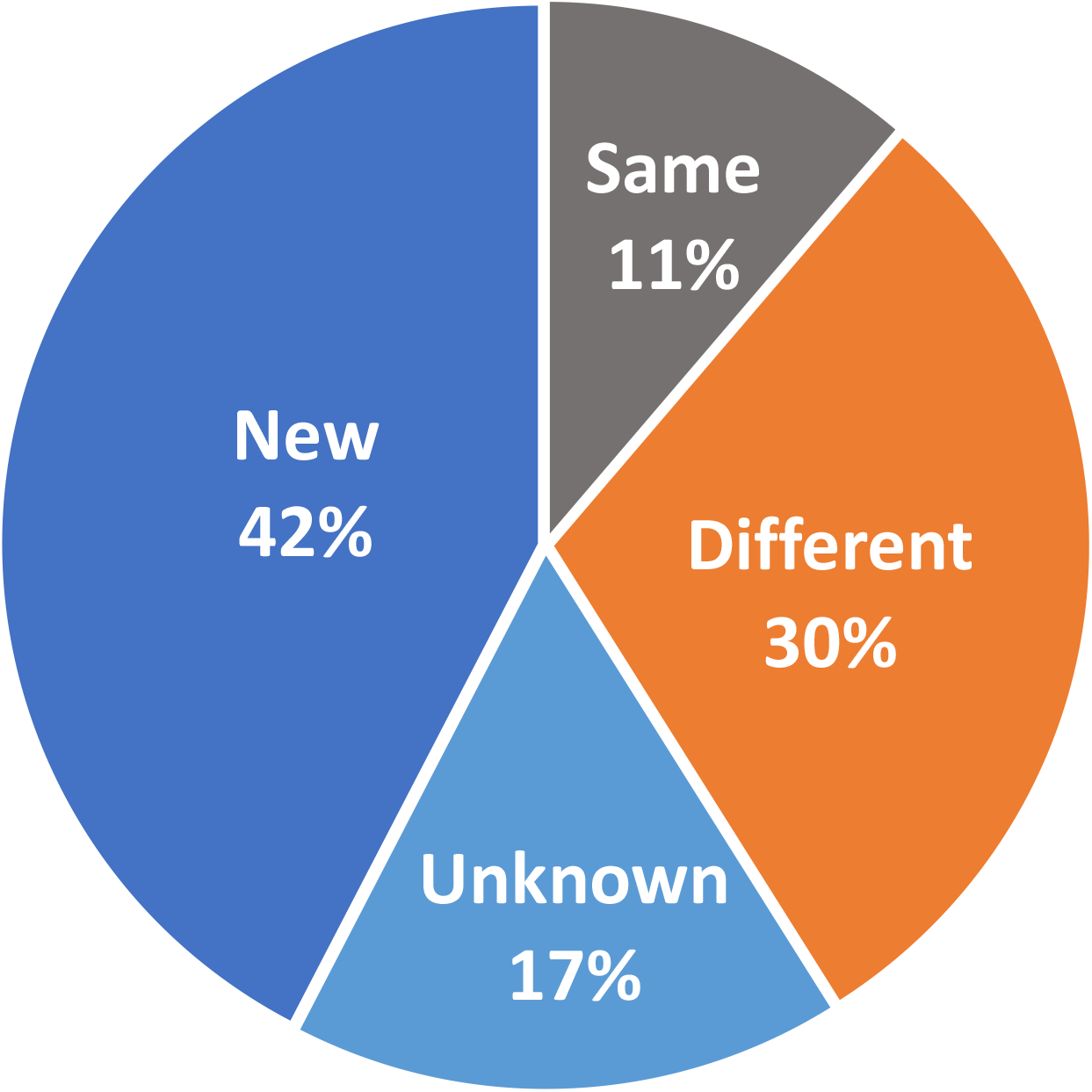
Total interactions identified by Rescaled Bliss Independence (RBI) contrasted to previously published results. Over half (59%) of the total interactions examined were untested and are therefore “new” interactions (42%) or were tested in the original study, but the interaction type was not specified or classified by the original study, and are therefore “previously unknown” interactions (17%) now classified using RBI. Only 11% of interactions analyzed with RBI remained the same as previously published results; 30% resulted in a different interaction than previously reported.

Of the combinations that resulted in the same interaction type when applying both the original method described in the published studies and the RBI, 41% (*n*=7) were additive, 53% (*n*=9) were synergistic, and 6% (*n*=1) were antagonistic (Table 1). Among the interactions reclassified by the RBI, 71% (*n=*32) were previously published additive interactions, all of which were reassigned as antagonistic (Figure 3). The remaining interactions reclassified by RBI were synergistic and reclassified as antagonism (22%, *n*=10) or additive (7%, *n*=3). We found that 82% (*n*=32) of additive and 45% (*n*=10) of synergistic net interactions were reclassified as antagonistic net interactions using RBI (Table 1). No previously identified interaction type (*n*=62) was newly re-classified as synergy, but 41% (*n*=9) of interactions reported as synergy were confirmed using RBI (Table 1, Figure 3).

**Table 1.** Comparison counts for each net interaction type originally reported in previously published results and how they were re-classified using Rescaled Bliss Independence (RBI). “Inconclusive” interactions under “Previously Published Results” correspond to cases for which no explicit interaction type is reported or investigated.

| Net Interactions |  | Rescaled Bliss Independence (RBI) |  |  |
| --- | --- | --- | --- | --- |
|  |  | Synergy | Additive | Antagonism |
| Previously Published Results | Synergy | 9 | 3 | 10 |
|  | Additive | 0 | 7 | 32 |
|  | Antagonism | 0 | 0 | 1 |
|  | Inconclusive | 18 | 10 | 61 |

**Figure 3.**
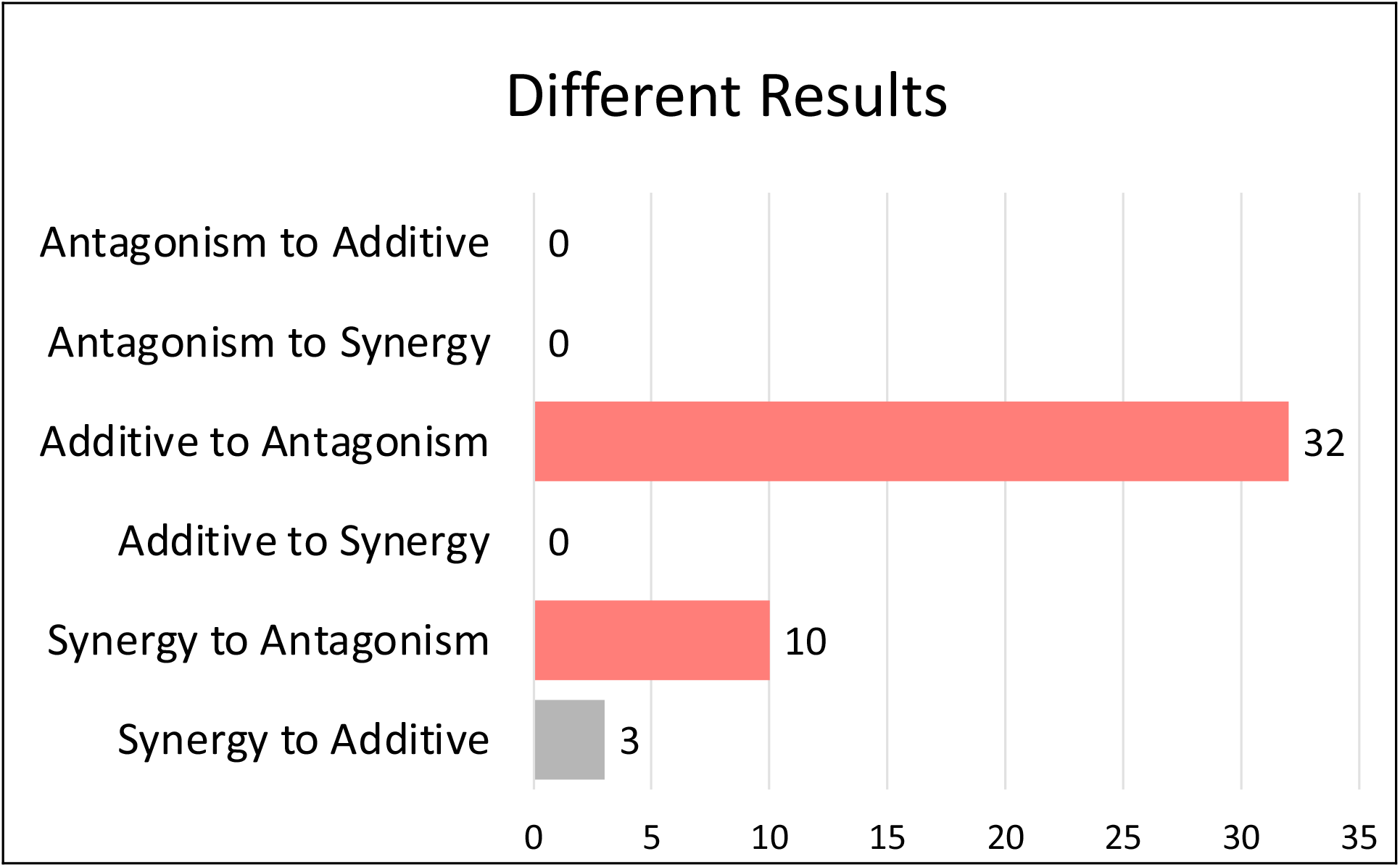
Traditional methods have difficulty identifying antagonistic interactions. Each bar shows the number of combinations that demonstrated a change from the previously published interaction type to the type of interaction with RBI.

We also examined the frequency of interaction types among net and emergent three-stressor combinations (Figure 4). Of the net interactions identified, we found that antagonism was the most prevalent interaction type at 69% (*n*=104). Of the antagonistic interactions, 34% (*n*=35) were suppressive. The remaining net properties were composed of 18% (*n*=27) synergistic and 13% (*n*=20) additive interactions (Figure 4A). Of the emergent interactions, we found that synergy and additivity were the leading interaction types across emergent properties–accounting for 47% (*n*=71) and 40% (*n*=61), respectively (Figure 4B). Antagonism accounted for 13% (*n*=19) of emergent interactions (Figure 4B).

**Figure 4.**
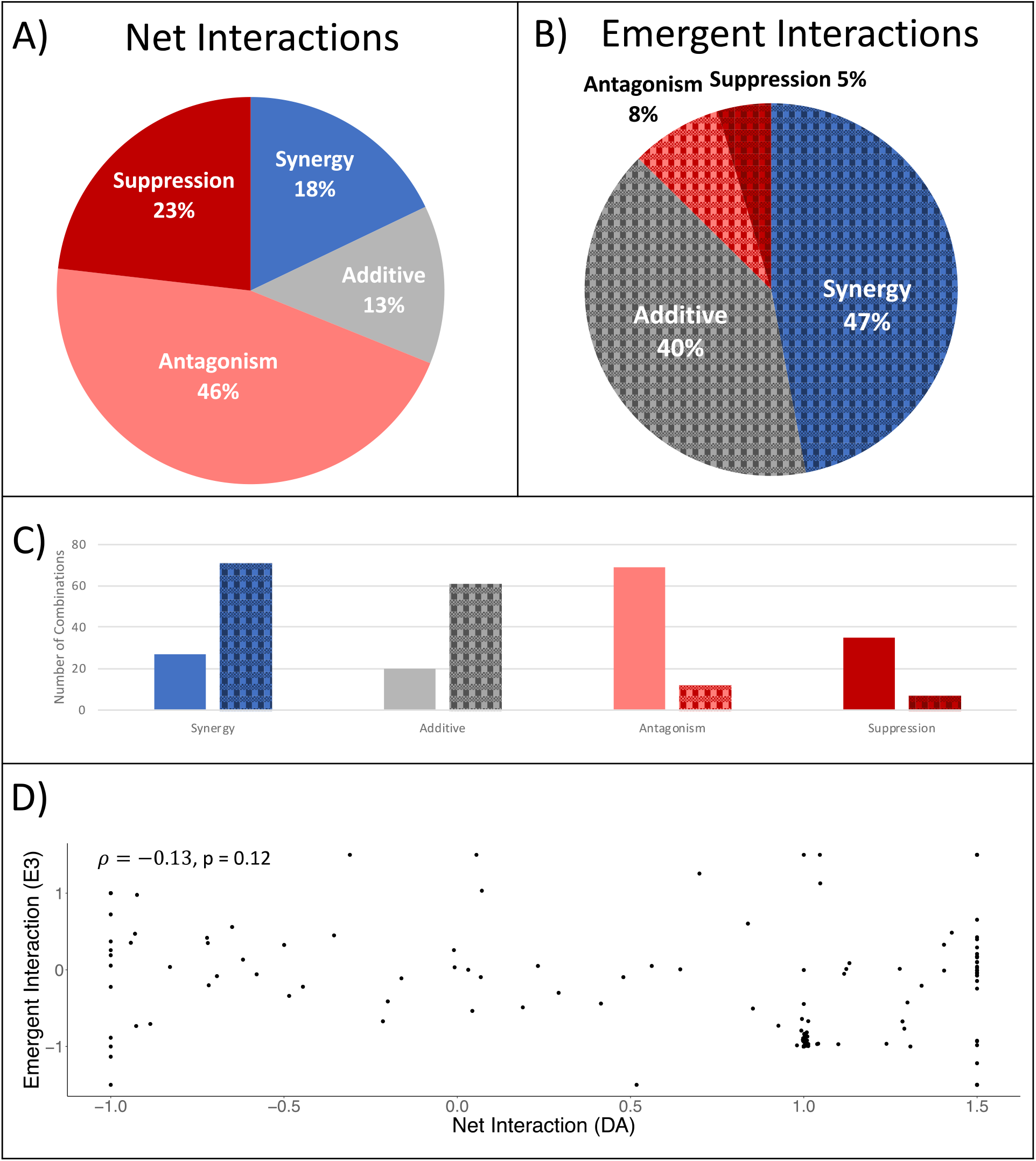
The composition of the net and emergent interactions using the RBI method. In each panel, gray represents additivity, blue represents synergism, and red represents antagonism. A darkening red illustrates an intensifying antagonism (e.g., antagonism → suppression). The plaid pattern represents emergent interactions, while solid colors represent net interactions. Panels A) and B) demonstrate the composition of the net and emergent interactions, respectively. Panel C) shows a direct comparison between the net and emergent interactions for each interaction type (synergy, additive, antagonism, buffering, and suppression). Panel D) shows no significant correlation between the net and emergent interaction values (Spearman correlation, *ρ =* −0.13, p = 0.12).

When comparing the frequency of interaction types among the net and emergent properties, we found that there were over three times as many instances of additivity in emergent interactions (emergent: *n=*61, net: *n*=20) (Figure 4C). There were also substantially fewer instances of emergent antagonistic interactions (*n*=19), including fewer suppressive interactions (*n*=7) among emergent when compared to net properties (*n*=104 total antagonism, including *n*=35 suppressive interactions). We then examined if there was a correlation between the net and emergent interactions. We did not find a significant correlation after performing a Spearman’s correlation (p = 0.12) (Figure 4D). We also observed more synergistic emergent interactions (emergent: *n=*71) than synergistic net interactions (net: *n*=27). Across all combinations, 65% (*n*=98) were found to have instances of “hidden suppression” where a pairwise combination is suppressed by the presence of a third stressor (e.g., better fitness with three stressors than with two stressors for negative stressor combinations). These interactions are only present in 3+ stressor interaction combinations. The comparison of the interactions’ distributions can be seen in Figure 5.

**Figure 5.**
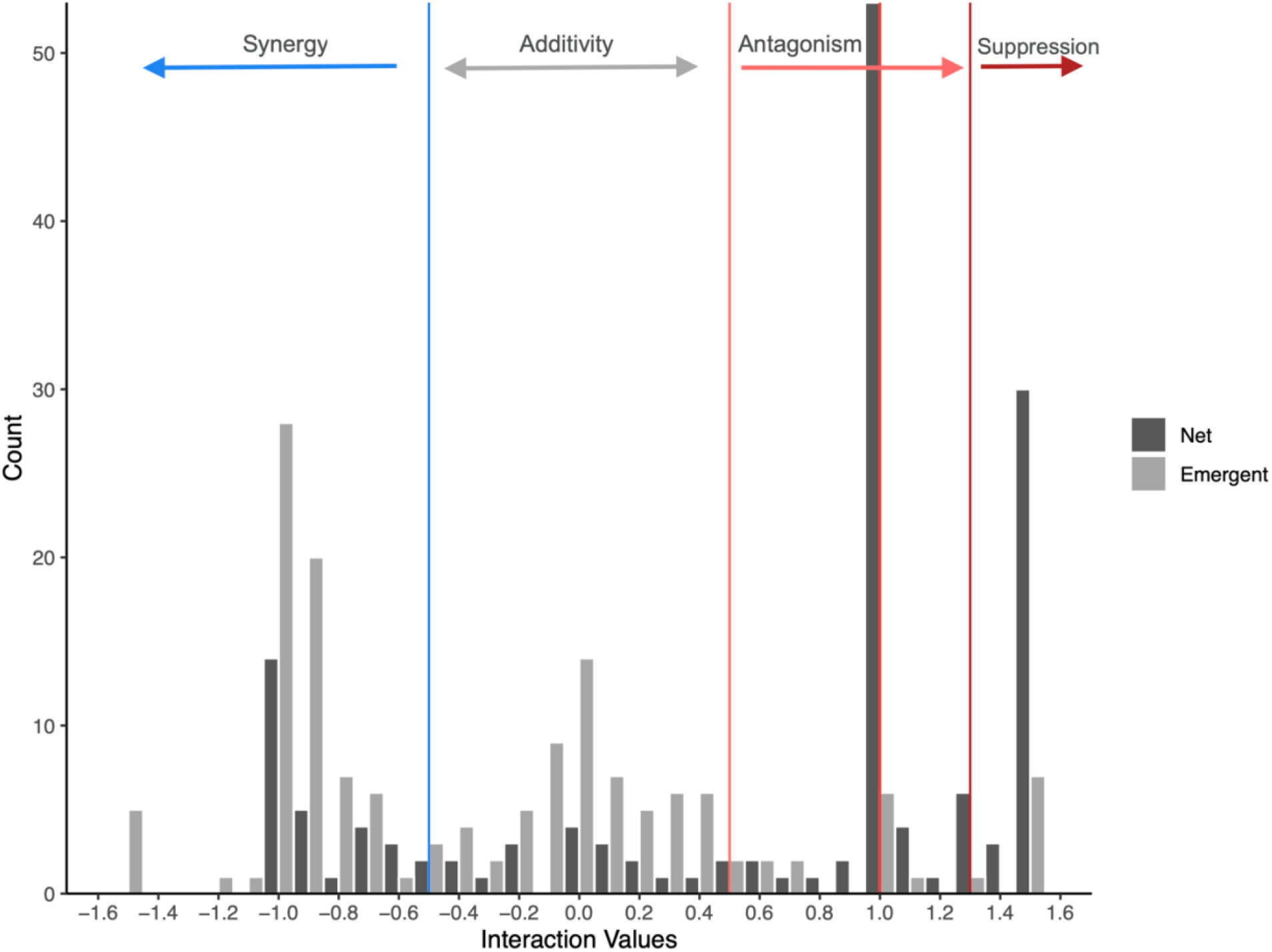
The distribution of interaction values of both net and emergent interactions. The distribution of net (DA) and emergent (E3) values. Cut-off values for each interaction type are as followed: synergy is less than -0.5; additivity is between -0.5 and 0.5; antagonism is above 0.5, suppression is above 1.3. Rescaled values are distributed across multimodal peaks. Thus, rescaling aids in the identification of interaction types through the use of these peaks.

## Discussion

In this study, we surveyed ecological literature published between 1920-2020 that examined the effect of three stressors simultaneously on population mortality, survival, or growth. After re-analyzing data from previously published results using a newly introduced framework, the RBI, we identified 151 interactions. We found that only 11% of the results generated by RBI matched those in the original studies, meaning that nearly 90% of interactions were classified as new (either unspecified or not investigated by the previous authors) or different interaction types (Figure 2).

Our results show that methods used in the original studies may have difficulty identifying antagonistic interactions (Figure 3). When comparing our findings using the RBI framework to those of the original findings, we found that interactions were more often reclassified as antagonisms than synergy, and antagonisms made up most interaction types (nearly 69%). Furthermore, there was only one instance of both RBI and original methods classifying combinations as antagonistic—although crucially, the one case that met the criteria of this study where a three-stressor interaction was previously classified as “antagonism.” In one example, an interaction was classified as synergistic using restricted maximum likelihood ANOVA methods when assessing the combined effects of UV-radiation, water temperature, and salinity stress on mollusk embryos (Przeslawski *et al*. 2005). However, during our reanalysis, we reclassified the interaction as antagonistic. Similar results were observed when using RBI to reanalyze work on insecticide combinations in anuran species (Boone 2008), pesticide combinations and food limitation in *Daphnia magna* (Shahid *et al*. 2019), and a combination of abiotic and biotic stressors in a seagrass (*Zostera noltei*) (Vieira *et al*. 2020). Traditional ANOVA and log-logistic methods initially classified these combinations as synergy, but, using RBI, they were reclassified as antagonism.

Higher-order interactions involving three or more stressors and emergent properties that arise from higher-order combinations are still poorly understood (Beppler *et al*., 2016; Tekin *et al*., 2018). We asked how often emergent properties appear in higher-order ecological stressor interactions. We find that emergent effects occur nearly 60% of the time, suggesting that emergent properties are common among higher-order ecological interactions. Moreover, we demonstrate that the RBI framework can identify higher-order emergent interactions that are overlooked or not explicitly explored when using traditional methods. For example, when using a general linearized model (GLM) to investigate the effect of pH, temperature, and oxygen availability on moon jellyfish, additive interactions were reported (Algueró-Muñiz *et al*. 2016). However, using RBI, we did not confirm the original authors’ conclusion and instead found antagonistic net interactions, and we also identified synergistic emergent interactions.

Our data also demonstrate that emergent properties persist across all interaction types—synergy, additivity, and antagonism. We demonstrated that synergy and additivity are the leading effects across emergent interactions—accounting for 47% and 40% of the total interaction types identified in our study, respectively (Figure 4B). Among the 13% of emergent antagonistic interactions we identified by RBI, 37% of them were characterized as suppressive (where one stressor reverses another stressor’s effects) (Figure 4).

In contrast to our findings, some prior studies observed more synergistic net interactions in comparison to antagonistic interactions when examining three-stressor interactions, even if antagonism was more common in two-stressor combinations (Crain *et al*. 2008; Maher *et al*. 2019). Our study examined over three times as many interactions as Crain *et al*. (2008) (*n*=151 and *n*=48 respectively). Maher and colleagues (Maher *et al*. 2019) found that synergies dominate three-stressor interactions in the coral microbiome using GLM and LMM models rather than RBI, which could explain the finding of synergistic interactions. This study also focused on a different biological scale—the microbiome *community* rather than a unique population’s fitness.

On the other hand, one major three-stressor interaction study, which observed the impact of low food, thermal stress, and elevated toxin levels on *Daphnia* populations, demonstrated that most interactions were antagonistic (Folt *et al*. 1999), similar to what we found here. Although emergent properties were not explicitly stated, the severity of the combined antagonistic effects differed from what would be predicted based on the sum of individual effects (Folt *et al*. 1999), suggesting that emergent properties played a role in population response. From a very different field, that of microbiology, antibiotic-combination studies reveal that higher-order emergent interactions were most often antagonistic than synergistic (Beppler *et al*. 2016; Tekin *et al*. 2018a). By identifying emergent interaction types, we can determine the combined effects of specific factors on populations in complex habitats which are subjected to multiple stressors at any given time.

Although our study addresses three-stressor interactions, our results are comparable to the previously mentioned two-stressor interaction studies in that additivity and antagonism were also found to be the most prevalent interaction types in a reanalysis of two-way interactions using RBI from ecological studies within the past 25 years (Tekin *et al*. 2020). In the two-stressor studies, 41% (*n*=286 of 840) of interactions were identified as additivity and 40% (*n*=278 of 840) as antagonism (Tekin *et al*. 2020). Interestingly, those results correspond well with previous reports that were not using RBI (Darling & Cote 2008; Côté *et al*. 2016; Jackson *et al*. 2016). This provides support for the idea that synergy may be overemphasized in the literature and that antagonism may occur more often than previously thought (Darling & Cote 2008), at least for net interactions. Synergy has been overemphasized in other biological disciplines, including research on antibiotic resistance (Singh & Yeh 2017).

Historically, whether three-stressor interactions exist and, if they do, to what extent they affect natural populations and ecosystems has been a subject of debate since the 1960s (e.g., Vandermeer 1969; Pomerantz 1981; Abrams 1983; Billick & Case 1994). At the population level, one major limitation in understanding these interactions is determining an applicable and generalizable model. By applying RBI, we were able to properly assess three-stressor interactions and determine that not only do emergent properties exist across biological systems but that they are also relatively common. Thus, a population’s response cannot necessarily be predicted by assuming additivity across stressors.

Further work still needs to be done to scale from population dynamics to community and ecosystem functioning across time (Côté *et al*. 2016; Brooks & Crowe 2019; Jackson *et al*. 2021). Beyond the scale of single-species populations, interactions between species and resources (Coyte *et al*. 2015; Butler & O’Dwyer 2020) and higher-order interactions between species (Kelsic *et al*. 2015; Grilli *et al*. 2017) have been shown to be important in modeling stability in ecological communities. Additionally, evolution in response to multiple stressor interactions and the fitness landscapes they form could influence adaptive dynamics (Ogbunugafor *et al*. 2016), population outcomes (Venturelli *et al*. 2015), and therefore broader eco-evolutionary dynamics. Indeed, understanding selection can help determine trajectories populations may take in adapting to given stressors (e.g., Toprak *et al*. 2012).

The implications of finding higher-order interactions extend beyond basic science. There is growing awareness that stressor interactions are crucial for population management and response predictions across systems. Pairwise interactions have received plenty of attention and have been the subject of many studies. In contrast, higher-order interactions and emergent interactions in 3+ stressor systems—which almost certainly present a more accurate representation of what natural populations face and will continue to face—remain less understood. Properly identifying these interactions is critical for managing ecological stressors (Brown *et al*. 2013; Piggott *et al*. 2015). The finding that the majority of emergent interactions are synergistic or additive rather than antagonistic, while the majority of combinations also exhibit hidden suppression, suggests that we need to identify *which* stressors are involved in a given system and what the impact may be if a given stressor is removed or decreased.

Crucially, identifying emergent properties can reveal hidden suppressive interactions (i.e., suppressive interactions that only occur among higher-order interactions). These hidden suppressive interactions could be particularly important for the ecological management of at-risk populations. In a three-stressor combination, the addition of a third stressor may suppress a two-stressor interaction. For example, when examining the combined effects of acidification, drought, and warming, the interaction between drought and acidification is suppressed by the effect of higher temperature on plankton producer biomass resulting in more biomass with all three stressors than with two (Christensen *et al*. 2006). Such an observation may be important for incorporating necessary mitigation strategies. In this example, alleviating acidification would result in lower biomass because it would undo the suppression of the interaction between warming and drought.

Instead, mitigating warming and drought would be a better strategy if the goal were to increase biomass. If the stressor interactions are not clearly understood or identified, the wrong stressor could be mitigated. These hidden suppressive interactions are common: the majority (65%) of our re-analyzed stressor combinations revealed hidden suppression between a pairwise interaction and a third stressor.

Throughout the ecological literature, investigations of three-stressor interactions involving biotic and environmental stressors remain scarce. We completed a thorough literature search of thousands of relevant results over the last 100 years and found that only 0.3% of research articles examining three ecological stressors qualified for our meta-analysis. Comparatively, when exploring pairwise interactions across the ecological literature, Tekin *et al*. (2020) found that nearly 8% of search results were applicable. Nevertheless, over the last decade, studies investigating higher-order interactions across disciplines have increased dramatically (Figure S2). Most of the studies that qualified for re-analysis here occurred within the past five years. The recent increase in higher-order interaction studies highlights how crucial it is that we extend our research beyond pairwise interactions to more accurately examine the effect of stressors in combination.

In conclusion, we show that this new RBI framework can be generalized from pairwise interactions to three or more stressors to examine how multiple stressors interact. RBI can distinguish between net and emergent interactions, providing greater insight into complex biological systems. From a basic science perspective, predicting higher-order interactions is essential to understanding how the combined effects of multiple stressors interact and impact diverse biological systems. From a conservation perspective, multiple stressor interactions can influence the population size of species of concern. Thus, correctly characterizing multiple stressor interactions can be crucial for developing management strategies to mitigate biodiversity loss.

## Acknowledgements

We thank Carissa Reulbach, Jasmine Noh, Eber Rivera, Lin Zhou, Kayla Edmunds, Naomi Lopez, Michael Yu, and Lilia Chahine for help with data collection. For funding, we thank James F. McDonnell Complex Systems Scholar Award, US National Science Foundation (ID: 1254159, http://nsf.gov/), UCLA Faculty Career Development Award, Hellman Foundation Award, and NIH/National Center for Advancing Translational Science (NCATS) UCLA CTSI Grant Number UL1TR001881.

