## Supplementary Material for "Meta-analysis using new methods for three-stressor combinations reveal substantial higher-order interactions and emergent properties"

Mailing address: 621 Charles E. Young Drive South, Los Angeles, California 90095, U.S.A.

Statement of authorship: P.J.Y. and V.M.S. conceived of the project and designed the study. V.E. and A.K. collected data. N.A.L.-H. performed the analysis of the data. E.S.D., S. B., N.A.L.-H., and P.J.Y. interpreted the data and all authors wrote and revised the manuscript.

Data accessibility: Datasets are available in Dryad for peer-review access and will be made publicly available upon acceptance.
DOI: https://doi.org/10.5068/D1W10G

Short title: Higher-order Ecological Interactions
Keywords: environmental stressors, multiple stressors, synergy, antagonism, additivity
Type of article: Letter
Number of words in the abstract: 148
Number of words in the main text: 4986
Number of references: 61
Number of figures: 5
Number of text boxes: 0
Number of tables: 1
Number of Supporting Information figures: 2
Number of Supporting Information text boxes: 3
Number of Supporting Information tables: 1
Number of Supporting Information references: 55

**Text S1. A summary of the issues with using ANOVA (ANalysis Of VAriance) for classifying interaction types as outlined in Tekin *et al.* (2020).**

The limitations and potential false inferences of applying ANOVA to test for and classify interactions are fourfold. **First**, variance in response measures within and across all treatments is determined by the number of experimental replicates, which are often limited in stressor-combination studies. This then leads to poor estimates of higher-order moments (e.g., kurtosis and skewness), potentially leading to inaccurate results. **Second**, though hidden replication (e.g., assuming variance is constant across treatments) is often used to justify applying ANOVA in these scenarios, hidden replication rests on the assumption that there is *no interaction* between variables (Welham *et al.* 2014). Thus, employing hidden replication when testing for interaction often invalidates the findings themselves. Further, non-linear pairwise interactions require carefully chosen data transformations when the assumptions of ANOVA are otherwise not violated (Pomerantz 1981; Billick & Case 1994; Gotelli *et al.* 1999). It is particularly important to transform the underlying additive model to a multiplicative model when stressors have large effects on populations (Segrè *et al.* 2005; Tekin *et al.* 2018a). **Third**, ANOVA assumes Gaussian or parametric distributions when comparing variances between treatments. Therefore, a large number of replicates per treatment would be necessary to accurately assess the variance of non-normal distributions and subsequently reconstruct ANOVA based on the non-parametric null model. **Fourth**, ANOVA methods do not allow for rescaling. Rescaling or normalization, relative to control baselines (e.g., population fitness with no stressors present), is often necessary to classify interaction types because different interaction types may result in similar unscaled values. Additionally, rescaling results in multimodal distribution of interaction values, aiding interaction classifications (Figure 5) (Segrè *et al.* 2005; Tekin *et al.* 2018b, 2020).

**Text S2. Mathematics of Rescaling Bliss Independence with Multiple Stressors that Inhibit Growth.**

Once interaction values are calculated as described in the methods it may be hard to distinguish a cut-off value for each type of interaction. To do this, rescaling is needed to transform a unimodal distribution of the raw interaction scores to become trimodal, allowing for clearer distinctions of truly antagonistic and synergistic interactions. For this study, we followed protocols developed by Tekin *et al.* (2016) to rescale. For both net and emergent synergistic interactions, we normalized to a lethal case because when measuring growth rates, relative fitness cannot be below zero.

$$DA_{rescaled} = \frac{DA}{|\mathbf{0} - w_A w_B|} \qquad DA_{rescaled} = \frac{DA}{|\mathbf{0} - w_A w_B w_C|}$$

When rescaling occurs for non-synergistic net interactions the interaction value is normalized to the minimum of the single stressor effects.

$$DA_{rescaled} = \frac{DA}{|\min(w_A, w_B) - w_A w_B|} \qquad DA_{rescaled} = \frac{DA}{|\min(w_A, w_B, w_C) - w_A w_B w_C|}$$

When rescaling occurs for non-synergistic emergent interactions we chose to normalize the interaction value to the minimum of the pairwise interactions. Tekin *et al.* (2016) recommend this normalization option because it may be more biologically relevant than other options.

$$E3_{rescaled} = \frac{E3}{|\min(w_A DA_{BC}, w_C DA_{AC}, w_C DA_{AB}) - w_A DA_{BC} - w_B, DA_{AC} - w_C DA_{AB}|}$$

**Text S3. Mathematics of Rescaling Bliss Independence with Stressors That Increase Growth**

For combinations that only had stressors that increased growth, we adapted the protocols developed by Beppler *et al.* (2016) and Tekin *et al.* (2016) to rescale. For the initial net (DA) and emergent (E3) interactions, the signs were reversed to keep synergistic interactions as negative values and antagonistic interactions as positive values following the equations below.

$$DA = w_A w_B w_C - w_{ABC}$$

$$E3 = w_{AB} w_C + w_{AC} w_B + w_{BC} w_A - 2 w_A w_B w_C - w_{ABC}$$

When rescaling synergistic interactions, the interaction value is normalized to the maximum positive value because there is no upper limit to what a synergistic combination of promoters can be. Ideally this maximal value would be infinity, however this is not practical. In an attempt to estimate this maximal value, we used twice the amount of the relative fitness of the highest-order combination.

$$DA_{rescaled} = \frac{DA}{|2w_{AB} - w_A w_B|} \qquad DA_{rescaled} = \frac{DA}{|2w_{ABC} - w_A w_B w_C|}$$

When rescaling occurs for non-synergistic net interactions, the interaction value is normalized to the maximum of the single stressor effects. This was done to keep with the same definition of rescaling to buffering, as described in Tekin *et al.* (2016): buffering normalizes to the most extreme of the single stressors.

$$DA_{rescaled} = \frac{DA}{|\max(w_A, w_B) - w_A w_B|} \qquad DA_{rescaled} = \frac{DA}{|\max(w_A, w_B, w_C) - w_A w_B w_C|}$$

When rescaling occurs for non-synergistic emergent interactions we chose to normalize the interaction value to the maximum of the pairwise interactions. Tekin *et al.* (2016) recommend this normalization option because it may be more biologically relevant than other options. Again, we chose to use the maximum of the single and pairwise interactions following the definition of buffering that normalizes to the most extreme effect of the single stressors and lower-order combinations that interact additively with the third stressor (Tekin *et al.*, 2016). When all stressors inhibit growth, the most extreme effect would result in the minimum amount of growth. Whereas, in combinations of stressors that each increased growth, the most extreme effect would result in the maximum amount of growth.

$$E3_{rescaled} = \frac{E3}{|\max(w_A DA_{BC}, w_C DA_{AC}, w_C DA_{AB}) - w_A DA_{BC} - w_B, DA_{AC} - w_C DA_{AB}|}$$

**Table S1.** Research articles included in our study. The author(s), publication year, habitat type, species, unique stressors, response variable, net interactions reported, and interactions identified by RBI are provided, including the emergent and net interactions. Growth responses vary in "units reported" as some are growth rates determined by change in population size or mass divided by experimental days ("growth rate"). Other growth responses are raw population size or mass at the conclusion of the experiment ("growth"). When reporting interactions, we list the type of interaction followed by the number of instances for that interaction type in that study (interaction type: number of instances).

| habitat | source | species | stressors | responses (units reported) | interaction reported (net) | interaction by RBI (emergent) | interaction by RBI (net) |
|---|---|---|---|---|---|---|---|
| **Estuary** | (Gobler *et al.* 2018) | *Menidia berylina* | $pCO_2$, temperature, food limitation | growth (mm), survival (%) | Synergy: 2 | Additive: 2 | Antagonism: 2 |
| **Freshwater** | (Boone 2008) | *Rana clamitans* | carbaryl, malathion, permethrin | growth (g) | Synergy: 1 | Synergy: 1 | Additive: 1 |
| | (Buck *et al.* 2012) | *Rana cascadae* | carbaryl, *Batrachochytrium dendrobatidis*, *Pseudacris regilla* | growth rate (mg/d) | Additive: 1 | Additive: 1 | Antagonism: 1 |
| | (Chen *et al.* 2008) | *Simocephalus vetulus* | triclopyr, pH, food availability | survival (%) | Additive: 1 | Additive: 1 | Antagonism: 1 |
| | (Chen *et al.* 2004) | *Simocephalus vetulus* | herbicide, pH, food availability | survival (%) | None: 2 | Synergy: 2 | Antagonism: 2 |
| | (Davis *et al.* 2018) | *Agapetus fuscipes*, *Silo pallipes* | sediment level, phosphorus, nitrogen | growth (individuals/mesocosm) | None: 6 | Additive: 1, Antagonism: 3, Synergy: 2 | Antagonism: 6 |
| | (De Coninck *et al.* 2013) | *Daphnia magna* | parasites, carbaryl, carbaryl pre-sensitivity | survival (%) | Synergy: 1 | Additive: 1 | Additive: 1 |
| | (Dinh *et al.* 2016) | *Coenagrion puella* | temperature, food limitation, chlorpyrifos | growth rate (ln(final mass)-ln(initial mass))/days) | Synergy: 1 | Additive: 1 | Synergy: 1 |

| | | | | | | | |
|---|---|---|---|---|---|---|---|
| | (Hasenbein *et al.* 2018) | *Hyalella azteca* | salinity, temperature, bifenthrin | survival (%) | None: 6 | Additive: 5, Antagonism: 1 | Antagonism: 3, Synergy: 3 |
| | (Hatch & Blaustein 2000) | *Rana cascadae* | pH, nitrate, UVB | survival (%) | Additive: 4 | Additive: 4 | Additive: 2, Antagonism: 2 |
| | (Hintz *et al.* 2019) | *Physella acuta, Helisoma trivolvis* | nutrients, predator presence, non-invasive snails | growth (g) | Additive: 1, None: 1 | Additive: 1, Synergy: 1 | Antagonism: 2 |
| | (Houde *et al.* 2019) | *Oncorhynchus tshawytscha* | salinity, temperature, hypoxia | survival (count) | None: 2 | Additive: 2 | Antagonism: 2 |
| | (Jansen *et al.* 2011) | *Daphnia magna* | carbaryl, *Pasteuria ramosa*, predation threat | survival (%) | Additive: 1 | Synergy: 1 | Antagonism: 1 |
| | (Janssens & Stoks 2013) | *Ischnura elegans* | food limitation, chlorpyrifos, temperature | growth rate ((ln(final mass)-ln(initial mass))/days) | Additive: 2 | Additive: 2 | Additive: 2 |
| | (Manzi *et al.* 2020) | *Daphnia longispina* x *galeata* hybrids | temperature, low food quality, parasite infection | growth rate (per capita rate of increase per day) | None: 2 | Additive: 2 | Additive: 1, Antagonism: 1 |
| | (Op de Beeck *et al.* 2018) | *Ischnura elegans* | temperature, CPF, density | survival (proportion intact dead larvae per mesocosm) | None: 2 | Synergy: 2 | Antagonism: 2 |
| | (Piggott *et al.* 2015) | *Gomphonema minutum, Encyonema minutum, Fragilaria vaucheriae, Gomphoneis minuta* var. *cassiae* | temperature, sediment, nutrients | growth (cells per $cm^2$ x1000) | None: 13 | Additive: 5, Antagonism: 1, Synergy: 7 | Antagonism: 13 |
| | (Relyea 2006) | *Rana clamitans* | predator presence, high pH, high carbaryl | growth rate (mg/d) | Additive: 1 | Additive: 1 | Antagonism: 1 |
| | (Shahid *et al.* 2019) | *Daphnia magna* | food limitation, Esfenvalerate, Prochloraz | survival (%) | Synergy: 5, None: 5 | Additive: 4, Antagonism: 4, Synergy: 2 | Antagonism: 1, Synergy: 9 |

| | | | | | | | |
|---|---|---|---|---|---|---|---|
| **Marine** | (Alguero-Muniz *et al.* 2016) | *Aurelia aurita* | pH, temperature, oxygen availability | survival (%) | Additive: 10 | Synergy: 10 | Antagonism: 10 |
| | (Andrew *et al.* 2019) | *Phaeocystis antarctica* | temperature, light, iron | growth rate (μ) | None: 1 | Additive: 1 | Additive: 1 |
| | (Blake & Duffy 2016) | *Zostera marina* | shade, temperature, grazers | growth (g) | Additive: 2 | Additive: 2 | Additive: 1, Antagonism: 1 |
| | (Büscher *et al.* 2017) | *Lophelia pertusa* | elevated $CO_2$, temperature, low food availability | survival (% per day) | Antagonism: 1 | Synergy: 1 | Antagonism: 1 |
| | (Castro-Sanguino *et al.* 2017) | *Halimeda heteromorpha* | light exposure, herbivory, nutrient enrichment | growth (new segment count) | None: 4 | Synergy: 4 | Antagonism: 4 |
| | (Dineshram *et al.* 2016) | *Crassostrea gigas* | temperature, reduced salinity, pH | survival (%) | Additive: 1 | Additive: 1 | Additive: 1 |
| | (Gamain *et al.* 2018) | *Zostera noltei* | temperature, pesticide mixture, copper | growth rate ((ln(final biomass)-ln(initial biomass))/days) | None: 1 | Additive: 1 | Synergy: 1 |
| | (Gil *et al.* 2016) | *Porites rus* | nutrient enrichment, sedimentation, overfishing | survival (% live tissue cover per colony) | Additive: 2 | Synergy: 2 | Antagonism: 2 |
| | (Gobler *et al.* 2018) | *Menidia beryllina* | diet, $pCO_2$, temperature | survival (%) | Synergy: 1 | Additive: 1 | Synergy: 1 |
| | (Hoadley *et al.* 2016) | *Symbiodinium trenchii* | temperature, nutrients, $pCO_s$ | growth (cells $cm^{-2}$) | Additive: 1 | Additive: 1 | Antagonism: 1 |
| | (Maulvault *et al.* 2019) | *Diplodus sargus* | temperature, triclosan exposure, acidification | growth (cm) | None: 1 | Additive: 1 | Additive: 1 |
| | (Oliver *et al.* 2019) | *Crassostrea gigas* | Imidacloprid, handling, air exposure | survival (%) | Additive: 1 | Synergy: 1 | Antagonism: 1 |
| | (Przeslawski *et al.* 2005) | *Dolabrifera brazieri, Bembicium nanum, Siphonaria denticulata* | temperature, salinity, light | survival (proportion) | None: 16, Synergy: 8 | Synergy: 24 | Antagonism: 24 |
| | (Kriegisch *et al.* 2019) | *Ecklonia radiata* | nutrient, sediment, density | growth (% cover) | Additive: 1 | Synergy: 1 | Antagonism: 1 |

| | | | | | | | |
|---|---|---|---|---|---|---|---|
| | (Vasquez *et al.* 2015b) | *Limulus polyphemus* | temperature, salinity, oxygen | survival (%) | None: 8 | Additive: 6, Antagonism: 2 | Additive: 1, Antagonism: 4, Synergy: 3 |
| | (Vasquez *et al.* 2015a) | *Limulus polyphemus* | temperature, oxygen, $H_2S$ | survival (%) | None: 1 | Additive: 1 | Additive: 1 |
| | (Vasquez *et al.* 2017) | *Limulus polyphemus* | temperature, salinity, oxygen | survival (%) | None: 12 | Additive: 9, Antagonism: 3 | Additive: 3, Antagonism: 5, Synergy: 4 |
| | (Vieira *et al.* 2020) | *Zostera noltei* | nutrients, sediment, density | growth (shoot density per corer) | Synergy: 1 | Synergy: 1 | Additive: 1 |
| **Terrestrial** | (Bednarska & Laskowski 2009) | *Pterostichus oblongopunctatus* | temperature, Chlorpyrifos, nickel | survival (%) | Synergy: 2 | Antagonism: 1, Synergy: 1 | Synergy: 2 |
| | (Dyer *et al.* 2003) | *Spodoptera frugiperda* | Piplartine, 4`-desmethylpiplartine, cenocladamide | growth (mg) | None: 1 | Synergy: 1 | Synergy: 1 |
| | (Janssens *et al.* 2017) | *Lestes viridis* | egg temperature, larval temperature, previous esfenvalerate concentration | survival (%) | None: 1 | Additive: 1 | Antagonism: 1 |
| | (Morgado *et al.* 2016) | *Porcellionides pruinosus* | Chlopyrifos, Mancozeb, soil moisture | survival (%) | None: 2 | Additive: 1, Synergy: 1 | Additive: 1, Synergy: 1 |
| | (Relyea 2006) | *Rana catesbeiana*, *Rana clamitans* | predator presence, high pH, carbaryl | survival (%) | Additive: 2 | Additive: 1, Synergy: 1 | Antagonism: 2 |
| | (Stevens & Gowing 2014) | *Anthoxanthum odoratum* | clipping, *Plantago lanceolata*, *Prunella vulgaris* | growth (g) | Additive: 1 | Antagonism: 1 | Additive: 1 |
| | (Ward *et al.* 1995) | *Meleagris gallopavo* f. *domestica* | dietary copper, water copper, Coccidiosis infection | survival (%) | Additive: 4 | Antagonism: 2, Synergy: 2 | Antagonism: 4 |
| | (Wilsey 1996) | *Stipa occidentalis* | $CO_2$, clipping, urea treatment | growth (g/plot) | Additive: 3 | Antagonism: 1, Synergy: 2 | Additive: 1, Antagonism: 2 |
| | (Wong *et al.* 2015) | *Spartina maritima* | nutrient availability, inundation, soil type | survival (%) | None: 1 | Synergy: 1 | Synergy: 1 |

### Supplementary Figures

#### PRISMA 2009 Flow Diagram

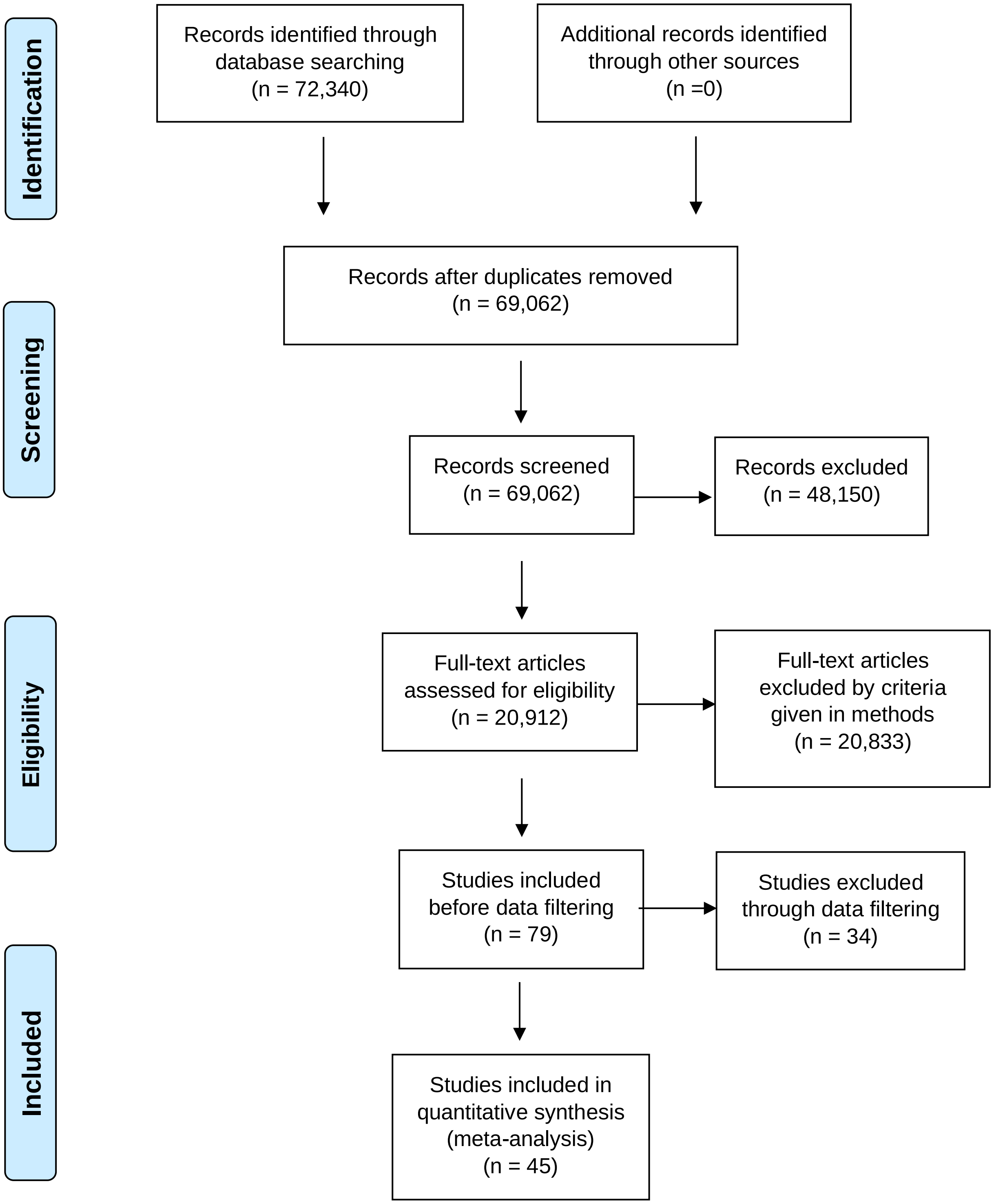

**Figure S1. PRISMA (Preferred Reporting Items for Systematic Reviews and Meta-Analyses) (Moher *et al.* 2009) Flow Diagram**. Using the *Web of Science* database, 45 out of 20,912 studies (records) were identified and included in our meta-analysis, resulting in 151 unique interactions.

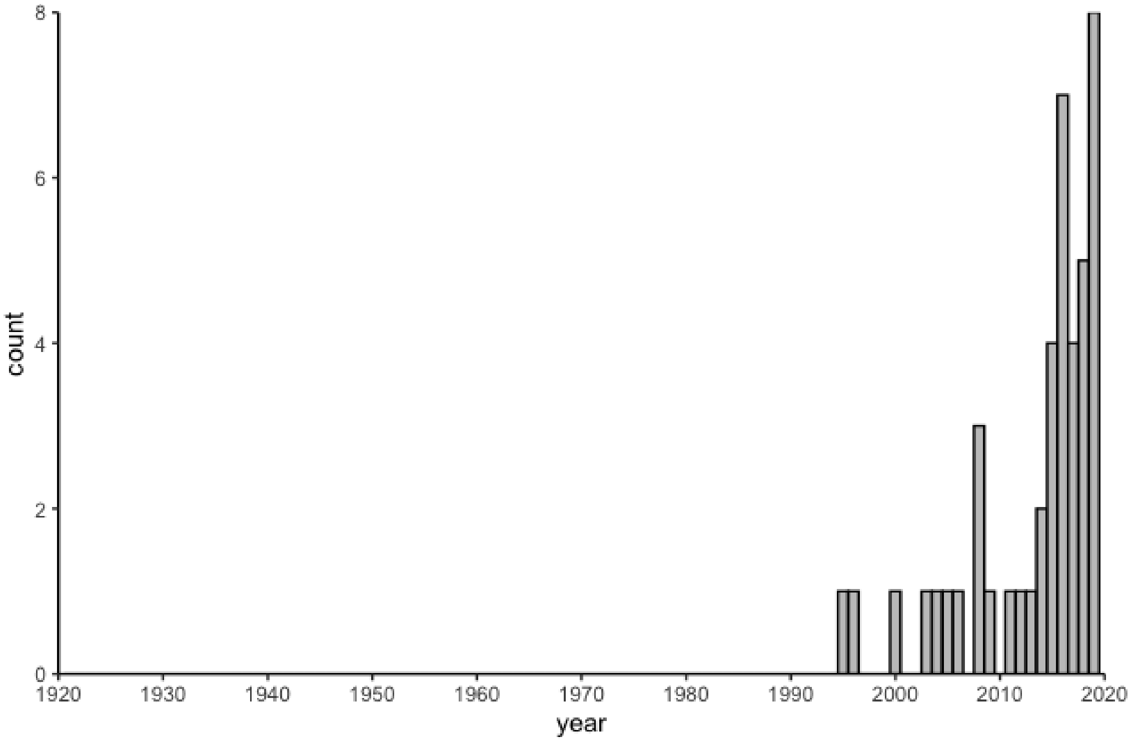


**Figure S2. The number of qualifying three-stressor studies that examined growth, mortality, or survival at the population level between January 1920-November 2020, by year.** Across a 100-year timespan, we identified 45 unique papers that were conducted in a factorial design that fit our data quality requirements needed for RBI. Most qualifying studies were conducted from 2016 to 2020 (*n*=24).
